# Degradation of non-methylated DNA by MsPJI impairs its usefulness as epigenetic tool

**DOI:** 10.1101/072090

**Authors:** María Belén Jerez, Maximiliano Juri Ayub

## Abstract

DNA treatment with sensitive and/or dependent restriction enzymes, followed by PCR amplification is a widely used approach for testing CpG methylation. Recently, MspJI has been characterized as a promisory tool for epigenetic analyses. In the present report, we describe that MspJI shows significant activity against non-methylated DNA, suggesting that additional caution and improvements would be required before applying this enzyme as a routine epigenetic tool.

## INTRODUCTION

In the last years, research on the epigenetic DNA modification has increased exponentially. In particular, knowing the methylation state of cytosine residues at CpG sites in animal genomes is a widely addressed issue, because 5-methylcytosine has been implied as a major regulatior of gene expression [1].

Two general basic approacches have been used for detecting cytosine methylation; namely bisulfite conversion and restriction with methylation-sensitive or dependent enzymes. The bisulfite based methods are powerful but cumbersome, since bisulfite modified DNA is unstable, containing many strand breaks, and difficult to amplify. Therefore, restriction based approcches are of choice for non-specialized laboratories. One of these approaches is based on the combination of genomic DNA restriction with PCR (preferably qPCR) amplification, using primers flanking the DNA region under study [2] (Figure 1). Recently, a novel and promisory methyl-dependent enzyme has been described: MspJI [3]. This enzyme has been characterized and described as cleaving the ^m^CNNR sequence, being inactive against non-methylated sequences. For this reason, it has been postulated as a powerful tool for epinegetic studies, being able to detect around 50% of methylated cytosines at CpG sites. Therefore, we tested this enzyme as a tool for the specific detection of cytosine methylation on mammalian genomic DNA.

**Figure 1.**
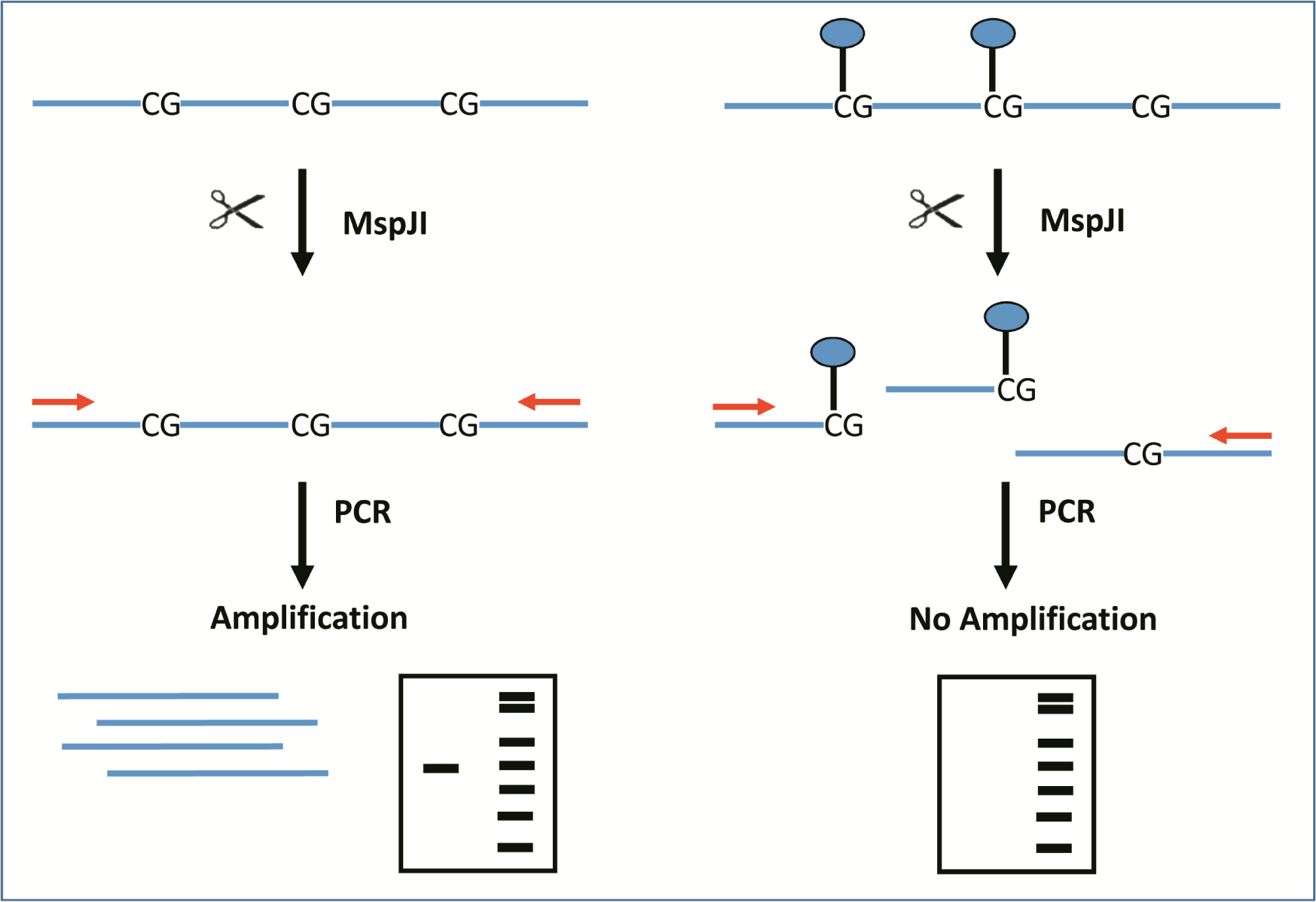
Schematic representation of the experimental design. Genomic DNA is treated with MspJI. If DNA molecules are non-methylated (left), template remains intact and can be amplified by PCR. If DNA is methylated (right), MspJI degrades template and weaker or no amplification take place. Partial methylation can be better detected by qPCR as a shift in Cq values.

## METHODS

### DNA purification

DNA was purified from mice peritoneal macrophages or *G. lamblia* axenic cultures using the Wizard® Genomic DNA Purification Kit (Promega), following the manufacturer instructions. Integrity of DNA was confirmed by agarose electrophoresis and quantification was carried out by absorbance at 260 nm using spectrophotometer (BioTek Instruments, Epoch™ /Take3™ Multi-Volume Plate).

### Digestion with MspJI

Genomic DNA (1µg) was digested following the manufacturer instructions, using 2 unit of MspJI (New England Biolabs) in the presence of 1 µl double-stranded DNA activator in a 30-µL volume. All reactions were incubated at 37 °C for 4 h or 8 h. The digestion products were visualized on 1% agarose electrophoresis.

### End-point PCR

PCR reactions were performed using 50 ng of genomics DNA incubated at 37°C in the presence or in the absence of MspJI. The following oligonucleotides were designed in order to amplify a 1,799 bp fragment from the IL-12b Fw: 5’ TCGGCCCCATATTGCTTTGT 3’ and Rev: 5’ ACAGCCTCTAGATGCAGGGA 3’. Thermocycling conditions were as follows; 1 cycle at 94°C for 30 sec; 35 cycles of 94°C for 30 sec, 63°C for 45 sec, 72°C for 2 min; followed by a final elongation step at 72°C for 10 min. Amplicons were visualized by electrophoresis on 1.5% agarose gel.

### Real-time PCR

PCR amplification mixtures contained 50 ng of genomic DNA, FastStart Universal SYBR Green Master (Rox) (Lifescience. Roche), 500 nM primers and UltraPure™ DNase/RNase-Free Distilled Water (Life technologies). Reactions were run on an ABI PRISM 7500 (Applied Biosystems). The cycling conditions comprised 2 min at 50°C, 10 min polymerase activation at 95°C and 40 cycles at 95°C for 15sec and 60°C for 60sec. The following primers were designed to amplify a 95bp CpG fragment of the beta actin gene: Fw CTTGATCTTCATGGTGCTAGGAG and Rev CAGTGCTGTCTGGTGGTAC. For PCR amplication of the 333 bp fragment from the IL-12b promoter, the following primers were used Fw: AAG TGT GTG GCT GGG AAG and Rev: GTT GAT GTT ACC TCCC TTCCTC.

## RESULTS AND DISCUSSION

**Figure 2** shows the digestion pattern after incubation of murine genomic DNA with MspJI. As expected, a smeared pattern was observed, consistent with the digestion of a densely methylated DNA. On the other hand, when genomic DNA of the protozoan *Giardia lamblia* showed a pattern similar to undigested sample. These results are consistent with a highly specific digestion of methyled DNA by MspJI, according to previous reports.

**Figure 2.**
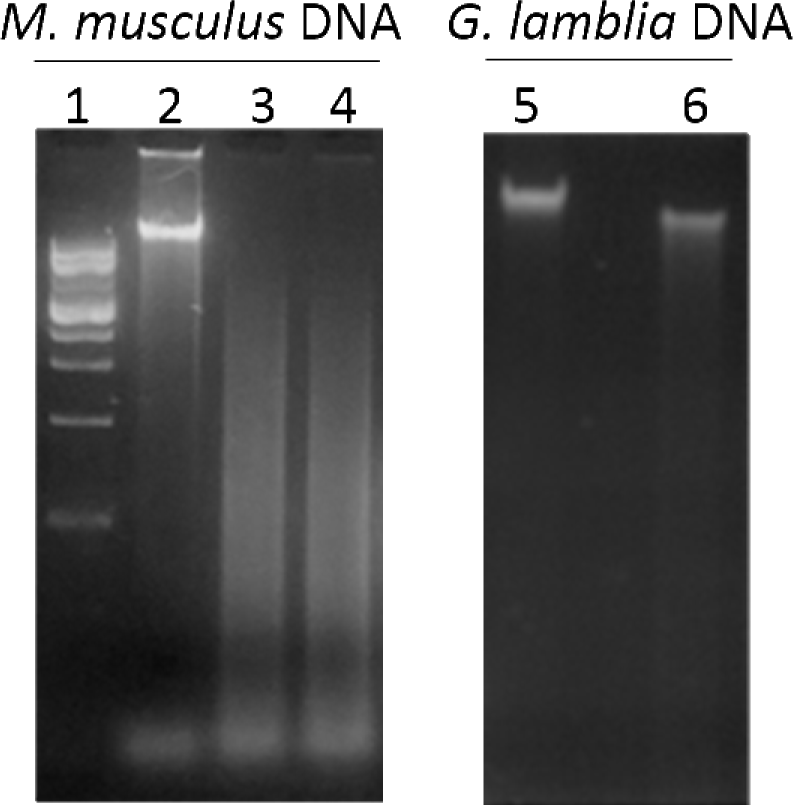
Electrophoresis of genomic DNA from *M. musculus* or *G. lamblia* digested with MspJI. Lane 1: 1kb DNA ladder; lane 2 and 5: undigested DNA; lanes 3 and 6: DNA incubated with MspJI for 4 hs; lane 4: DNA incubated with MspJI for 8 hs.

Next, we used MspJI digested mice DNA as template to amplify a 1,799 bp fragment from the IL-12b gene (GenBank: AH004859) (nt -1693bp to +106 bp, numbers according to the transcription initiation site), containing 39 CpG sites. Notably, digestion with MspJI completely prevented the amplification of the selected region, suggesting a high methylation state (**Figure 3**).

**Figure 3.**
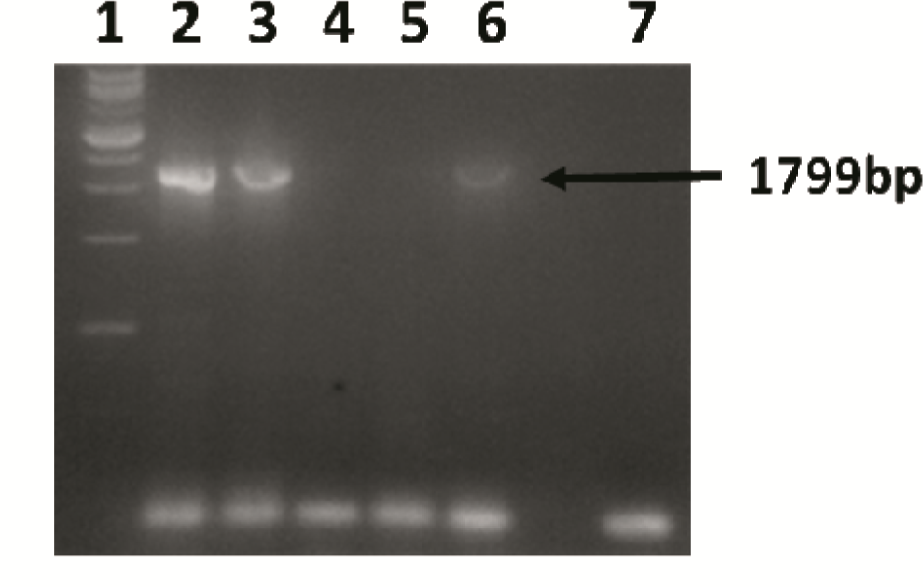
PCR amplification of IL-12b promoter gene. Genomic. DNA was incubated in the absence (lanes 2 and 3) or the presence (lanes 4 and 5) of MspJI, and used as template. Positive (lane 6) and negative (lane 7) amplification controls were analyzed in parallel.

This result prompted us to set a quantitative amplification analysis by using qPCR instead of end-point PCR. We selected two different PCR targets: a -333 bp sub fragment on the IL-12b promoter including 14 CpG sites (a CpG island; nt -1136 to -801, numbers according to the transcription initiation site) and a 95 bp fragment belonging to beta-actin gene (GenBank: NC_000071.6; nt 142904067 to 142904161, nt +999 to +1093 according to the transcription initiation site) lacking of CpG sites. Surprisingly, in both cases, enzyme treatment produced a 2-3 cycle’s shift of the Cq value (**Figure 4**), showing that DNA degradation was not dependent on CpG methylation.

Although this would be rather unusual, it can be postulated that methylated cytosines at sequences other than CpG were present in the genomic samples and recognized by MspJI. In fact, it has been reported non-CpG methylation in stem cells and, more recently, in differentiated mammalian cells [4]. Therefore, in order to evaluate the specificity of the assay using a *bona fide* non-methylated sample, we replaced genomic DNA template by a diluted PCR product. Notably, treatment of this non-methylated DNA sample with MspJI, PCR amplification was prevented, definitively confirming that MspJI degrades non-methylated DNA at a significant extension. These results demonstrate that MspJI is inadequate for epigenetic analyses based on methylation-dependent restriction coupled to qPCR strategy (**Figure 5**).

**Figure 4.**
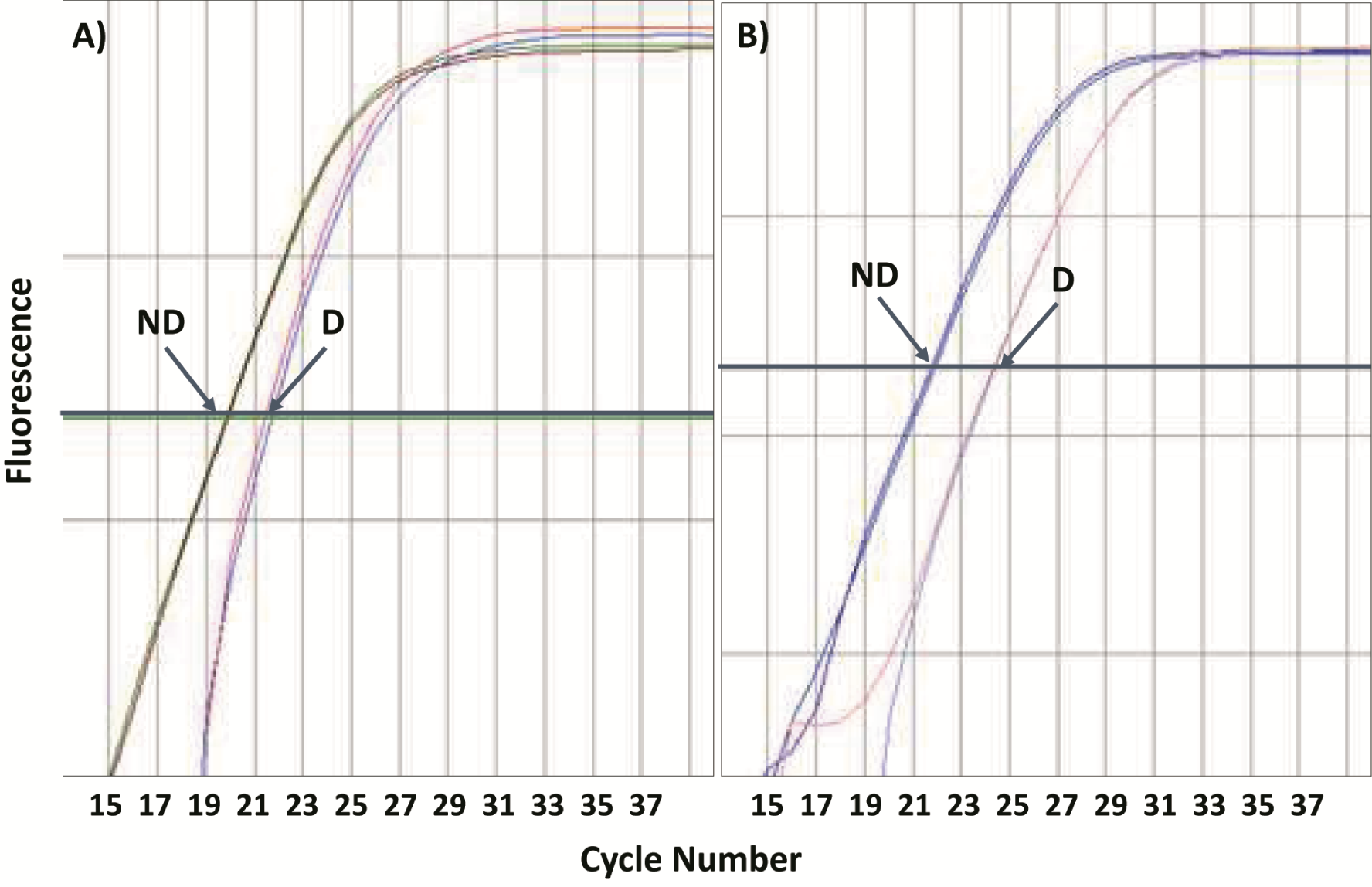
Effect of MspJI treatment on the real time amplification of genomic templates. qPCR reactions were performed using murine genomic DNA digested (D) or not digested (ND) with MspJI. The target sequences contained 14 (A) or none (B) CpG sites. The curves are representative of 12 independent experiments performed in duplicates.

**Figure 5.**
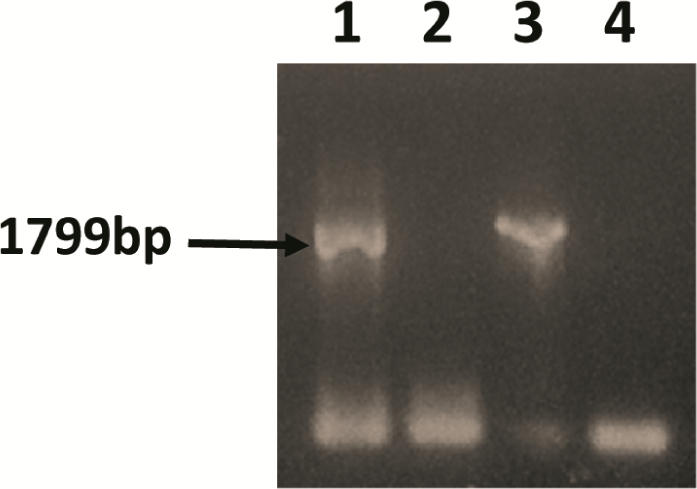
Effect of MspJI on the amplification of non-methylated DNA. PCR product of 1799 bp was incubated in the absence (lane 1) or presence (lane 2) of MspJI. Treated samples were re-amplifiied by PCR. Positive (lane 3) and negative (lane 4) controls were included.

Concluding, we have found that even when MspJI selectively degrade methylated DNA, its activity against non-methylated DNA is not negligible. Therefore, cautions should be taken when using this enzyme for epigenetic studies.

## ACKNOWLEDGMENTS

We are grateful to Dr. Maria L. Mascotti and Dr. Walter J. Lapadula for their helpful comments and criticisms about the manuscript.

